# De Novo Design of a Protein Binder to Probe Gas Channel and Enhance the Oxygen Tolerance of [NiFe] Hydrogenase

**DOI:** 10.1101/2025.11.19.689374

**Authors:** Xuan Sun, Wenjin Li, Wangzhe Li, Hang Luo, Qi Xiao, Leyan Zhang, Yilin Fan, Peiyu Jiang, Geng Wu, Liyun Zhang

## Abstract

[NiFe]-hydrogenases exhibit outstanding catalytic performance for H_2_ production and oxidation but suffer from extreme O_2_ sensitivity, limiting their biotechnological potential. The identity and functional roles of gas channels that connect to the buried [NiFe] active site have remained unclear, impeding the rational engineering of O_2_-tolerant enzyme variants. In this study, we present an artificial intelligence–guided de novo protein binder design strategy, utilizing compact protein binders as selective molecular probes to map hydrophobic O_2_ diffusion pathways in *Escherichia coli* hydrogenase-2 (Hyd-2). By integrating RFdiffusion, ProteinMPNN, and AlphaFold, approximately 100,000 candidate binders were computationally screened, leading to the identification of two high-affinity binders, L1 and L2. Structural, biophysical, and electrochemical analyses reveal that L1 specifically occludes the primary O_2_ ingress channel, thereby enhancing the enzyme’s oxygen tolerance by more than threefold, whereas L2, which targets a putative secondary channel, exerts a negligible effect, demonstrating the nonfunctionality of this pathway. These results provide the first direct experimental evidence for a hierarchical organization of O_2_ diffusion channels in [NiFe]-hydrogenases and establish binder-mediated channel occlusion as a generalizable, mutation-free approach for probing gas transport mechanisms and selectivity in metalloenzymes.

## Introduction

[NiFe]-hydrogenases are efficient and reversible catalysts for H_2_/H^+^ conversion, featuring a deeply buried bimetallic active site with three electron mediating iron-sulfur clusters. ^[1]^ Their exceptional activity highlights significant potential for biotechnological applications and the development of biomimetic hydrogen-evolving systems, ^[2]^ however, most of these enzymes exhibit oxygen sensitivity, which remains a major bottleneck. Molecular oxygen readily diffuses into the protein matrix, leading to the formation of inactive states and oxidative damage to the active site. ^[3]^ Over the past two decades, structural studies have proposed several models of gas diffusion networks in [NiFe]-hydrogenases. ^[4-6]^ Crystallographic analyses have revealed multiple hydrophobic channels connecting the protein surface to the buried active site, which are hypothesized to facilitate both H_2_ delivery and O_2_ intrusion. ^[5, 7]^ However, direct experimental evidence identifying the primary entry pathway remains lacking. Furthermore, the X-ray structure corresponds to a monomeric heterodimer composed of the large and small subunits, ^[6]^ whereas certain hydrogenases, such as Escherichia coli Hyd-2, exist as dimers of heterodimers in solution. ^[8]^ These discrepancies highlight the need for experimental validation to clarify the functional gas transport pathways and their roles in hydrogenase activity and inhibition. Although the crystal structure of Hyd-2 has been determined [8], it remains uncharacterized with respect to its oxygen diffusion channel. As a representative oxygen-sensitive [NiFe]-hydrogenase, it is still unknown whether Hyd-2 shares a conserved O_2_ access route with other oxygen-sensitive members of this enzyme class.

Several distinct strategies have been employed to enhance oxygen resistance, one of which involves blocking the gas channel to restrict oxygen access. For example, residues Val74 and Leu122 in Desulfovibrio fructosovorans [NiFe]-hydrogenase are positioned near the active site; the double mutants L122M-V74M and L122F-V74I exhibit increased oxygen tolerance due to bulky amino acid side chains that narrow the gas channel and limit O_2_ diffusion to the active site. ^[4]^ In addition to protein engineering, material-based strategies have been developed to protect hydrogenases from oxygen-induced inactivation. ^[9]^ Synthetic redox polymers, ^[10-12]^ peptide hydrogels, ^[13]^ and protein-derived shells can encapsulate enzymes ^[14]^, forming physical and redox-active barriers that scavenge or restrict O_2_. ^[10, 11]^ These external matrices preserve the intrinsic catalytic properties of the enzymes but are still limited by poor durability, low gas selectivity, and mass transfer resistance. Achieving higher specificity requires strategies that directly and selectively block O_2_ diffusion through defined gas channels. Such targeted occlusion not only enhances oxygen tolerance but also serves as a mechanistic probe to experimentally verify whether previously proposed diffusion pathways are indeed functional conduits for gas transport.

Recent breakthroughs in deep learning-based artificial intelligence (AI) have revolutionized protein and peptide design. ^[15]^ Structure prediction frameworks such as AlphaFold ^[16]^ and RoseTTAFold, ^[17]^ together with generative platforms like ProteinMPNN[18], RFdiffusion, ^[19]^ and DiffDock, ^[20]^ now enable the de novo creation of binders with experimentally validated binding and stabilizing functions. ^[21, 22]^ These approaches have already been applied to enhance enzyme stability and generate high-affinity scaffolds confirmed by structural studies ^[23]^. Yet, no study has leveraged AI to design binders that selectively block O_2_ diffusion. Here, we introduce a de novo protein design strategy to create binders that specifically bind to the openings of the gas channels of *E. coli* Hyd-2 to investigate to probe certain function of defined protein surface and engineering substance selectivity of enzyme, providing both a theoretical framework and a practical foundation for expanding the utility of hydrogenases. In this study, the binding proteins we designed specifically target the entrance region of the gas diffusion pathway and bind to the enzyme surface, forming a physical barrier that effectively restricts oxygen intrusion while preserving substrate mass transfer, thereby significantly enhancing the oxygen tolerance of Hyd-2. Furthermore, our experimental approach provides a means to validate the gas diffusion pathways predicted by Caver: by systematically assessing the blocking effects of distinct binding proteins on two putative entry sites and integrating experimental data on changes in oxygen tolerance, the primary oxygen diffusion pathways are confirmed through wet-lab experiments.

## Results and Discussion

The program Caver ^[24]^ and MOLE ^[25]^ were employed to predict potential O_2_ diffusion pathways, with residue R479 designated as the starting point (Figure 1a), based on the crystal structure of E. Coil. Hyd-2 (PDB ID: 6EN9). ^[8]^ Channel 1, characterized by a minimum diameter (dmin) of approximately 1.1 Å and a length of approximately 53 Å, is identified as the primary gas pathway, whereas channel 2 (dmin of approximately 1.1 Å and length of approximately 37 Å) may serve as a secondary route (Figure 1b). The analysis revealed two channels that traverse multiple subunits and converge prior to reaching the [NiFe] active site. These pathways are predominantly lined with conserved hydrophobic residues such as Val117, Leu179, Trp16, Phe161, Leu162 in the HybC subunit, creating a low-polarity environment conducive to O_2_ diffusion. ^[26]^ A similar gas channel has been proposed by the Scheerer group for the O_2_-tolerant membrane-bound [NiFe] hydrogenase (MBH) from Ralstonia eutropha. ^[5, 6]^ Comparable findings have also been reported in Caver-based analyses of the [NiFeSe] hydrogenase from Desulfovibrio vulgaris Hildenborough. ^[27]^

**Figure 1.**
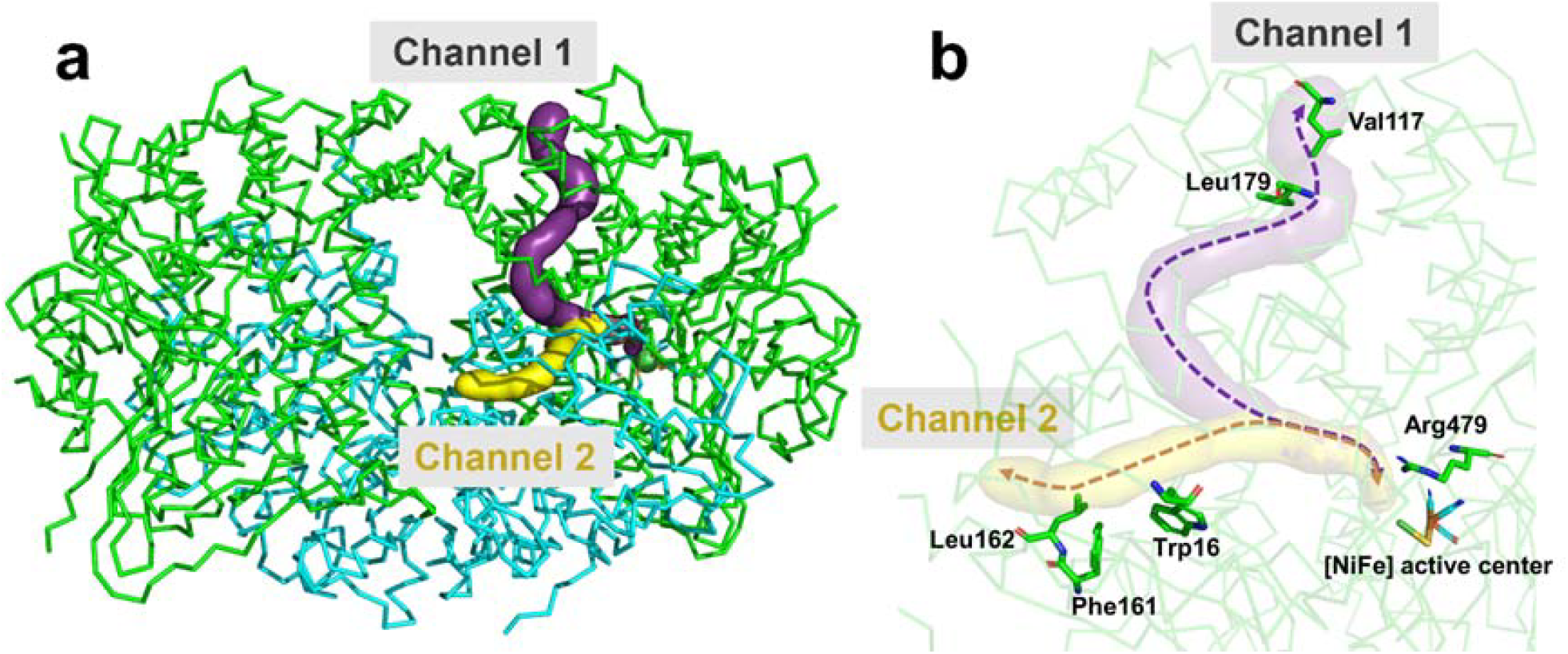
Predicted oxygen diffusion channels in Hyd-2. (a) Overall structure of Hyd-2 illustrating the predicted hydrophobic O_2_ diffusion pathways (purple and yellow) calculated using Caver and visualized in PyMOL. The large (green) and small (cyan) subunits of Hyd-2 (PDB ID: 6EN9) are shown in stick representation. (b) Surface representation of channel 1 (purple) and channel 2 (yellow), highlighting the [NiFe] active site and conserved channel-lining residues displayed as sticks.

Based on the aforementioned predictions and analyses of oxygen diffusion pathways, subsequent efforts will focus on designing binding proteins to block the entry points of these pathways, thereby restricting oxygen access to the interior of the enzyme molecule. We implemented a “block-and-lock” strategy in which small, designed protein binders sterically occlude channel entrances while maintaining the enzyme’s structural integrity and preserving native electron and proton transfer pathways. An iterative design–filter–selection pipeline was performed to progressively integrate generative modeling, structural fidelity assessments, and energetic evaluation, as shown in Figure 2a. Step 1: generative backbone design and sequence optimization. Using RFdiffusion, ^[19]^ we generated compact α-helical bundle topologies with protein length ranging from 70 to 140 residues, while enforcing a relatively flat, contiguous interaction interface complementary to the channel entrance (Figure S1). This process resulted in approximately 10,000 unique backbone structures. Each backbone was optimized with ProteinMPNN, ^[21]^ generating ∼100,000 sequences enriched for buried hydrophobic core residues to ensure conformational stability and polar interfacial residues predicted to form specific hydrogen-bond networks with Hyd-2. Step 2: Stability pre-filtering. Sequences unable to form stable hydrophobic cores in silico were discarded, reducing the pool to ∼30,000 candidates (Figure S2). Step 3: Complex modeling with AlphaFold2-multimer. The remaining designs were docked onto the HybO–HybC heterodimer of Hyd-2 (HybO and HybC are the large and small subunits of Hyd-2, respectively). Complexes with predicted alignment error (PAE) >8 or root mean square deviation (RMSD) >1 Å relative to the designed binding pose were removed, yielding 1,000 high-confidence binder–Hyd-2 complexes (Figure S3). Step 4: Fold verification with AlphaFold3. To confirm structural fidelity, each binder was predicted in isolation using AlphaFold3. ^[28]^ Designs with confidence scores (ipTM) > 0.9 were retained, narrowing the pool to 100 candidates (Figure S4). Step 5: Interaction selection. The binder-Hyd-2 interactions were evaluated by calculating the Rosetta interface energy, providing a quantitative thermodynamic analysis. A stringent cutoff of ΔG < −40 Rosetta Energy Units (REU), empirically correlated with nanomolar binding affinities, distilled the library to 10 lead binders (Figure S5). Overall, this process achieved a four-orders-of-magnitude down-selection, reducing the initial pool of approximately 100,000 sequences to five candidate pairs per target channel entrance, with integration of orthogonal criteria including stability, interface precision, and thermodynamic favorability.

**Figure 2.**
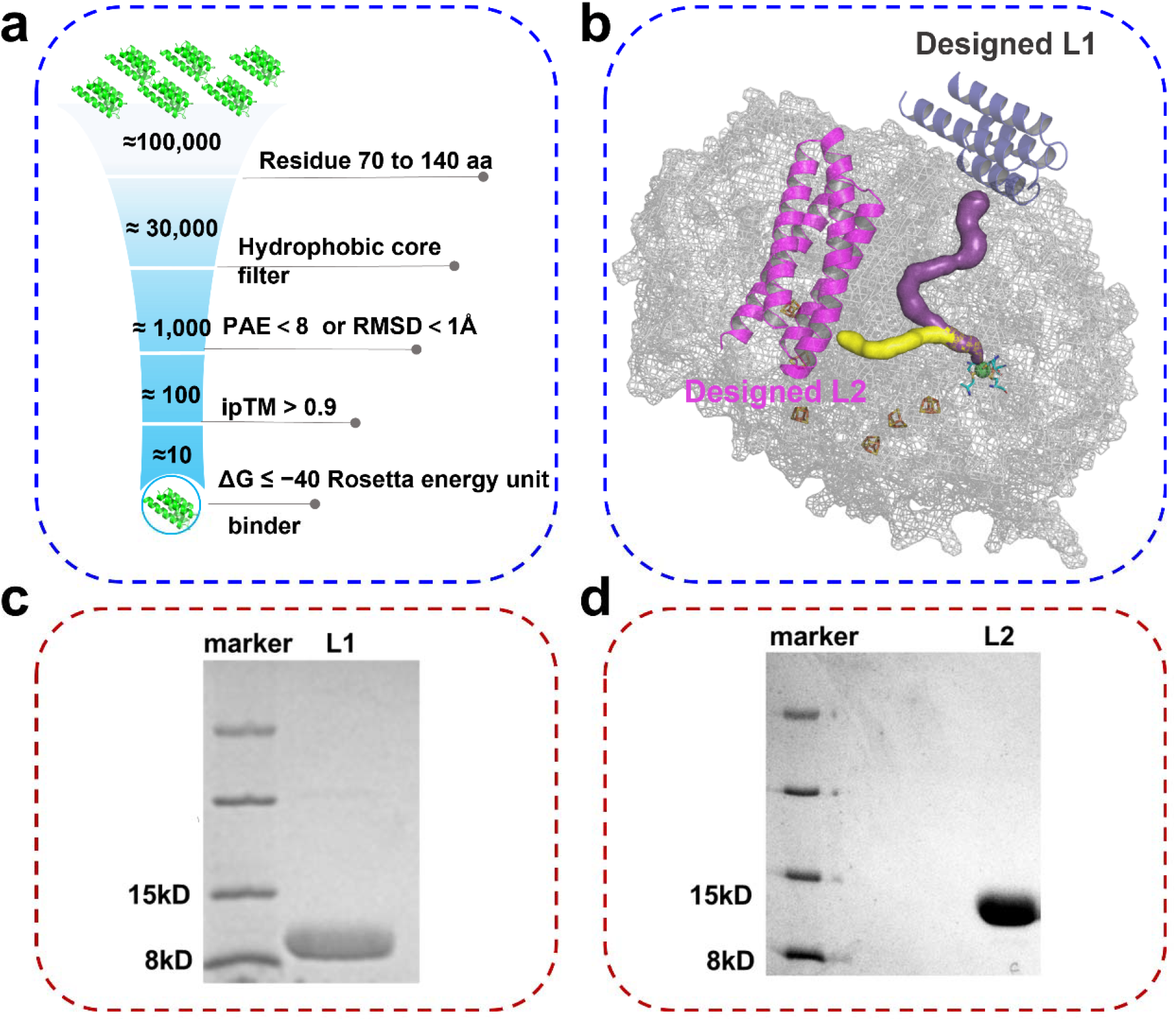
AI-guided de novo design and validation of protein binders blocking O_2_ diffusion channels in Hyd-2.(a) Workflow of the AI-guided de novo binder design and computational screening. Candidate sequences (70–140 residues) were generated and filtered based on hydrophobic core stability, structural confidence (predicted aligned error < 8 or backbone RMSD < 1 Å), and model quality (ipTM > 0.9). Final binders were selected according to the predicted binding affinity (ΔG ≤ −40 Rosetta energy units).(b) Representative structural model of Hyd-2 showing two designed binders (L1, grey; L2, magenta) positioned at the entrances of the gas channels (channel 1, purple; channel 2, yellow), sterically blocking O_2_ diffusion toward the [NiFe] active site.(c, d) SDS–PAGE analysis (18%) of purified binders L1 (8 kDa) and L2 (13 kDa).

Next, we prepared and evaluated the candidate binders using molecular biology techniques. The coding sequences of ten candidate protein binders were cloned into expression vectors, and the corresponding proteins were expressed in E. coli BL21 (DE3) cells via an inducible expression system. During the expression and purification process, candidates exhibiting low solubility or insufficient expression levels were excluded. Ultimately, two candidate binder pairs amenable to efficient production were successfully prepared and identified by SDS-PAGE, as shown in Figures 2c, 2d, S6, and S7, with their sequences provided in Table S1. Subsequently, these binders were individually co-incubated with the Hyd-2 enzyme to generate the Hyd-2+L complexes. The oxygen tolerance of these complexes was initially assessed through electrochemical cyclic voltammetry (CV, Figure S8-S12). Ultimately, the binders that conferred enhanced oxygen tolerance to Hyd-2— L1 bound to channel 1 and L2 bound to channel 2—were selected for further in-depth investigation (sequence information in bold is provided in Table S1). Structural modeling further confirmed precise shape complementarity and dense side-chain packing between the binders and their respective target channel entrances (Figure S13). Electrostatic surface analysis revealed strongly complementary charge distributions between L1 and channe 1 as well as between L2 and channel 2 (Figures S13 and S14), indicating high binding specificity. Both binders adopt tightly packed α-helical bundle folds stabilized by internal hydrophobic cores, consistent with aqueous solubility and conformational stability.

Surface plasmon resonance (SPR) was employed to characterize the direct binding interactions between the designed binders and E. coli Hyd-2. The resulting sensorgrams (black curves) exhibited excellent fit to a Langmuir 1:1 binding model (red curves) across all tested concentrations (Figures 3a and 3b), with no systematic deviations observed during the association or dissociation phases. These findings indicate specific and kinetically well-defined binding, with no significant contributions from surface heterogeneity or rebinding effects. The dissociation constant (Kd) of L1 was 112 nM, whereas that of L2 was 2.39 μM, demonstrating that both binders exhibit substantial affinity for Hyd-2, with L1 binding significantly more tightly than L2. These results confirm the rational design of the binders to achieve strong and specific interactions, thereby establishing a robust foundation for subsequent functional studies.

**Figure 3.**
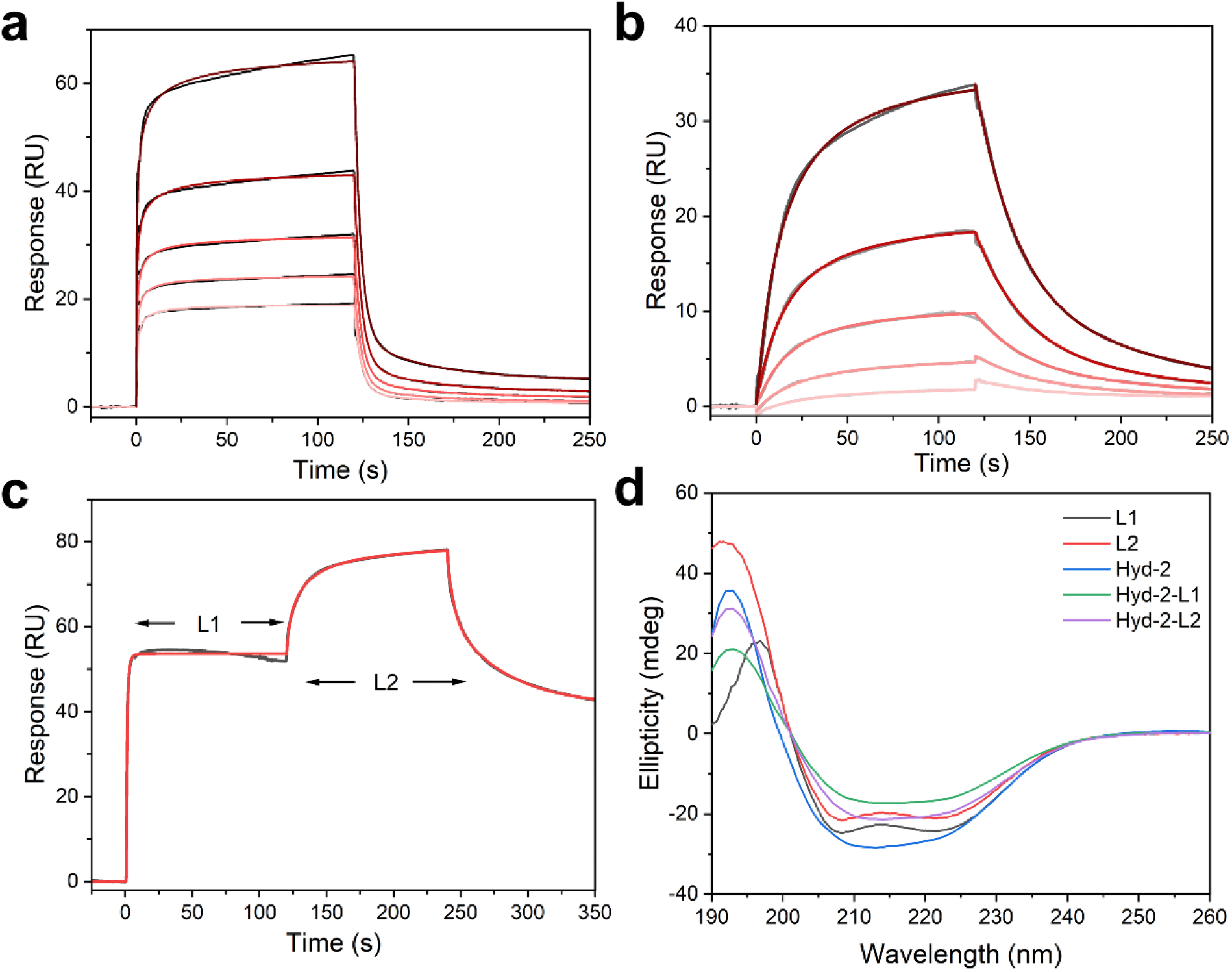
Biophysical characterization of binder binding to Hyd-2. SPR sensorgrams of the binding kinetics for the different concentrations of L1 (a) and L2 (b) with immobilized Hyd-2 (The red lines indicate the fitted data that was used to compute the binding affinities). c, SPR sensorgrams for investigating the different binding between the binders and immobilized Hyd-2. Data are presented as real-time graphs of response units (RUs) against time. d, Circular dichroism (CD) spectra of L1(black curve), L2(red curve), Hyd-2 (blue curve), and the complexes Hyd-2-L1 (green curve) or hyd-2-L2 (purple curve) in 0.1 M Tris-HCl buffer (pH 6.0). Measurements were performed at 25 °C.

This pair of binders was designed to selectively target distinct gas channels of Hyd-2. To validate this, a single-channel SPR competition assay was conducted. In the first injection, L1 was introduced, yielding a binding response of approximately 55 resonance units (RU, Figure 3c). A subsequent injection of L2 over the same channel resulted in an additional increase to approximately 80 RU, indicating further binder binding without displacement of L1. This stepwise increase suggests that L1 and L2 occupy non-overlapping binding sites on Hyd-2, ^[29, 30]^ consistent with their proposed roles in selectively modulating distinct O_2_ transport pathways. Notably, the addition of a sufficient amount of binder (Hyd-2: binder molar ratio of 1:5) to block both gas channels did not significantly alter the enzyme’s secondary structure (Figure 3d).

In the subsequent experiment, we employ direct electrochemical techniques, specifically protein film voltammetry, ^[31, 32]^ to conduct a detailed comparison of the effects of oxygen tolerance on Hyd-2. The cyclic voltammograms for the Hyd-2 (Figure 4a and 4b) were recorded at 10 mV s^-1^, during which, at approximately 0 mV on the forward sweep, 0.267 mL of O_2_-saturated buffer (25 °C, 1 atm, ∼ 1.2 mM) was injected into the 20 mL cell solution (final concentration ∼ 16 μM). The injection potential (at 0 V vs. SHE) was chosen such that O_2_ was not present in solution at potentials causing its reduction at the electrode. The voltammograms show immediately that O_2_ affects the activity of Hyd-2. The catalytic current drops very rapidly, indicating inhibition by O_2_. In contrast, the current drops by only 60 % when Hyd-2 is exposed to the same amount of O_2_ in presence of L1, establishing clearly that this enzyme continues to catalyze H_2_ oxidation. In stark contrast to L1, L2 exhibits a negligible effect on enhancing the oxygen tolerance of Hyd-2, indicating that blocking the so-called “channel 2” fails to effectively mitigate oxygen-induced inactivation. This suggests that “channel 2” does not function as a gas diffusion pathway. This finding is somewhat unexpected, as channel 2 exhibits a high degree of conservation in both oxygen-sensitive and oxygen-tolerant hydrogenases reported to date. ^[4,5.27,33]^ Nevertheless, this study represents the first direct experimental validation that this channel does not serve as the route for external O_2_ to enter Hyd-2.

**Figure 4.**
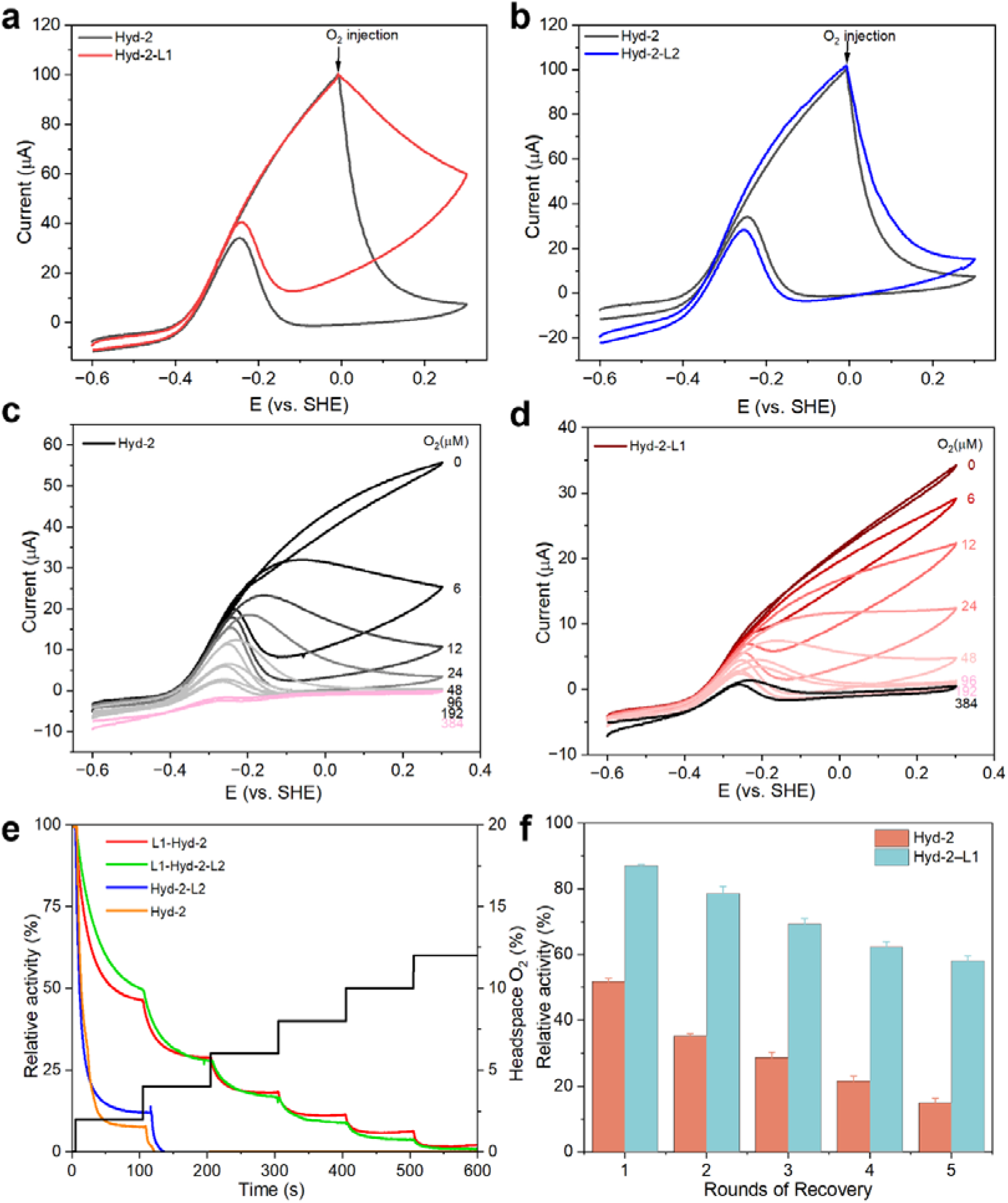
Biophysical characterization of binder binding to Hyd-2. SPR sensorgrams of the binding kinetics for the different concentrations of L1 (a) and L2 (b) with immobilized Hyd-2 (The red lines indicate the fitted data that was used to compute the binding affinities). c, SPR sensorgrams for investigating the different binding between the binders and immobilized Hyd-2. Data are presented as real-time graphs of response units (RUs) against time. d, Circular dichroism (CD) spectra of L1(black curve), L2(red curve), Hyd-2 (blue curve), and the complexes Hyd-2-L1 (green curve) or hyd-2-L2 (purple curve) in 0.1 M Tris-HCl buffer (pH 6.0). Measurements were performed at 25 °C.(e) Response of the H2 oxidation current to O2 exposure for Hyd-2 and Hyd-2–L1, measured by chronoamperometry at 0 V. Conditions: pH 6.0, 30 °C, ω = 3000 rpm, N2 carrier gas, total flow rate 1000 sccm. (f) Relative H2 uptake activity of Hyd-2 and Hyd-2–L1 during repeated cycles of deactivation and reactivation, derived from chronoamperometric measurements by deactivate with oxygen and +300 mV and recovery with hydrogen and −600mV.

Subsequently, enzyme activity was measured under varying O_2_ concentrations. As the O_2_ concentration increased from 6 to 384 μM, native Hyd-2 exhibited a rapid loss of activity, with an I_50_ (concentration causing 50% activity reduction) of 5 μM and complete inactivation occurring at 48 μM (Figure 4c). In contrast, The L1 bounded Hyd-2 shows much attenuated catalytic activity loses to O_2_ titration, resulting in an I_50_ of 17 μM, representing a more than three-fold improvement in oxygen tolerance (Figure 4d). In the chronoamperometric experiment, the enzymes were initially incubated under a constant atmosphere of 80% H_2_ and 20% N_2_ at a fixed potential of 0 V (vs. SHE), followed by exposure to 2%, 4%, 6%, 8%, 10%, and 12% O_2_ for 100 s. This exposure resulted in a rapid decrease in current for Hyd-2 to approximately 90% of its initial value, after which the current remained relatively stable for an additional 100 s. Notably, Hyd-2 was completely deactivated within approximately 120 seconds under a 4% O_2_ atmosphere (Figure 4e). In contrast, Hyd-2-L1 complex exhibited significant improvements in both the time to oxidative deactivation and the tolerance to higher oxygen concentrations. Research has demonstrated that the oxygen-induced inactivation of Hyd-2 is reversible. ^[34, 35]^ Upon gradual replacement of O_2_ in the headspace with H_2_, its oxidative activity progressively recovers. As shown in Figure 4f, Hyd-2 bound to L1 exhibits remarkable stability and functional performance throughout multiple cycles of oxygen inactivation and hydrogen reactivation. Collectively, these results demonstrate that L1 effectively blocks the major O_2_ pathway, thereby enhancing the oxygen tolerance of Hyd-2. This effect provides a potential strategy for preserving hydrogenase activity under aerobic conditions.

To well understand how the designed binders restrict O_2_ transport, we performed all-atom molecular dynamics (MD) simulations of Hyd-2 bound to L1, L2, or both binders simultaneously under explicit solvent conditions (400 nanoseconds per system, Figure S15). In the binder-free enzyme, O_2_ molecules readily diffused through two solvent-exposed channels and frequently approached within 3 Å of the [NiFe] active center (Figure 5a), consistent with previous structural models of oxygen ingress. ^[6]^ Binding of L1 at channel 1 creates a hydrophobic cavity that transiently traps O_2_ near the channel entrance, effectively blocking the diffusion pathway (Figure 5b). In our MD simulations, radial distribution function analysis shows that in Hyd-2–L1, O_2_ completely disappears within 3 Å of the active site, while a new density peak emerges at approximately 10 Å, indicating that L1 imposes a physical barrier to oxygen diffusion. MD simulations also revealed that O_2_ molecules were transiently trapped within the hydrophobic core of L1 through weak van der Waals contacts with surrounding nonpolar residues, forming a dynamic molecular “gatekeeper” that impeded O_2_ diffusion. Density and contact-frequency analyses identified a pronounced occupancy pocket and an associated free-energy minimum near the channel entrance, indicating a metastable, hydrophobically stabilized site that restricts O_2_ mobility without permanent binding (Figure 5b). L2 binding at channel 2 also restricted O_2_ diffusion but was less effective. While the frequency of O_2_ penetration was reduced, a residual inner-shell peak at ∼5 Å persisted, indicating that O_2_ could still reach the active site via the unblocked channel 1 (Figure 5c). When both L1 and L2 were present, O_2_ remained >10 Å away from the catalytic center throughout the simulation, demonstrating near-complete exclusion of oxygen. In both Hyd-2 and Hyd-2–L2, O_2_ frequently sampled the critical 3 Å region, resulting in rapid inactivation. In contrast, no O_2_ molecules reached within 5 Å in the Hyd-2–L1 complex, confirming that L1 blocks the primary oxygen channel.

**Figure 5.**
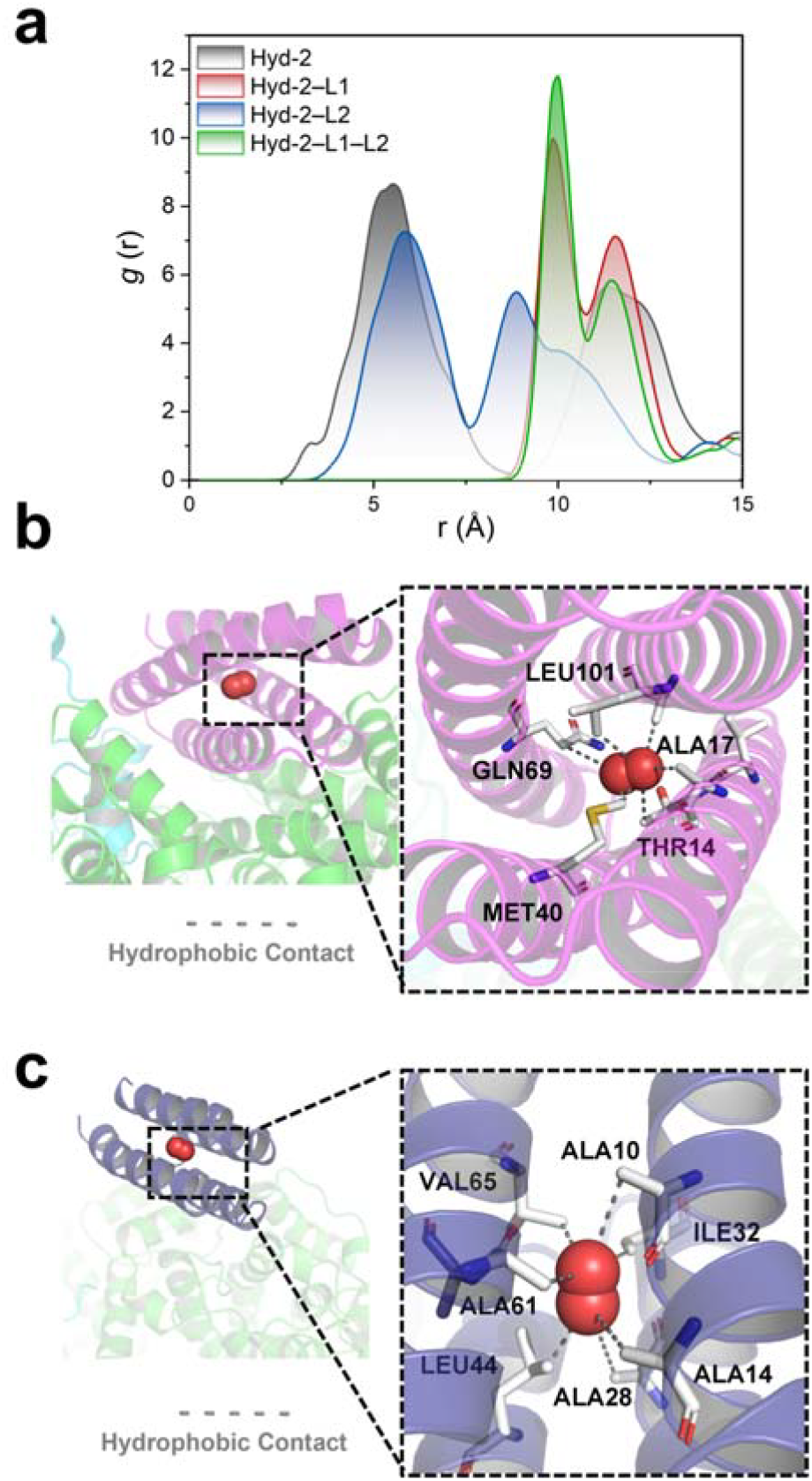
Molecular dynamics simulations revealing binder-mediated blocking of O_2_ access to the Hyd-2 active site. (a) Radial distribution function (RDF) profiles of O_2_ molecules at the entrances of channels 1 and 2 in different systems: apo Hyd-2 (black), Hyd-2–L1 (blue), Hyd-2– L2 (red), and Hyd-2–L1–L2 (green). (b, c) O_2_ trapping within the hydrophobic cores of binders L1 and L2, respectively. Insets highlight amino acid side chains constraining O_2_ through van der Waals and other noncovalent interactions.

Tracking the diffusion of gaseous substrates and inhibitors within proteins is crucial for elucidating enzyme function but remains experimentally elusive, particularly for transient species such as O_2_. As an oxygen-sensitive [NiFe]-hydrogenase, Hyd-2 provides an ideal model for dissecting oxygen transport pathways and improving oxygen tolerance. Here, we integrated molecular dynamics simulations with AI-guided *de novo* binder design to directly map and validate the O_2_ diffusion channels in Hyd-2. Unlike conventional “soak-and-freeze” ^[4, 5, 7]^ or mutagenesis-based strategies, ^[36]^ this design-driven approach enables experimental interrogation of gas channels without perturbing the buried catalytic core. Our findings unequivocally identify channel 1 as the primary conduit linking the protein surface to the [NiFe] active site. Although structurally conserved among [NiFe]-hydrogenases, ^[5]^ putative channel 2 does not contribute to effective O_2_ diffusion. This result may be attributed to the distinctive heterodimeric structure of Hyd-2. ^[8]^ This structural insight provides a mechanistic explanation for the long-standing challenge of enhancing oxygen tolerance in [NiFe]-hydrogenases and establishes a framework for rational engineering targeting key residues at channel entrances rather than global random mutagenesis.

From a methodological perspective, the external, shape-complementary binders act as molecular “caps” at solvent-exposed entrances, creating reversible barriers that restrict O_2_ influx while preserving H_2_ diffusion. This binder-mediated gating circumvents common mutagenesis pitfalls such as global conformational coupling or loss of catalytic turnover and offers a generalizable tool for testing channel hierarchy and selectivity. Functionally, the improved O_2_ tolerance observed upon channel blocking directly validates the predicted pathway and highlights the potential of using rationally designed binders to modulate gas transport in metalloenzymes.

Evolution has naturally equipped certain [NiFe]-hydrogenases with defensive adaptations—such as hydrophobic filtering channels, ^[37]^ SeCys substitution at the Ni site, ^[38, 39]^ or redox-active proximal [4Fe-3S] clusters ^[40]^—to mitigate O_2_ damage. Yet, most [NiFe]-hydrogenases remain highly sensitive to O_2_. The strategy presented here emulates and extends these natural mechanisms through *de novo* protein design, achieving selective gas gating via engineered molecular interfaces. Based on previous reports on material protection strategies ^[10-13]^ and the findings of this study, it is evident that reliance solely on physical barriers is insufficient to achieve complete enzyme protection. Future efforts to enhance the oxygen tolerance of hydrogenases should integrate a synergistic strategy that combines physical barrier functions with chemical scavenging mechanisms.

## Conclusion

In summary, this study establishes an AI-guided *de novo* binder design strategy as an effective means to experimentally probe and modulate gas diffusion pathways in [NiFe]-hydrogenases. By integrating molecular dynamics simulations with rational binder design, we directly identified channel 1 as the primary O_2_ transport route to the buried [NiFe] active site of E. coli Hyd-2, while excluding the functional contribution of the conserved channel 2. The designed binder L1 acts as a molecular cap that selectively blocks O_2_ ingress without disturbing catalytic integrity, thereby enhancing the enzyme’s oxygen tolerance by more than threefold. These findings provide the first direct experimental evidence linking channel architecture to O_2_ sensitivity in [NiFe]-hydrogenases and elucidate a mechanistic basis for their limited natural tolerance. Beyond revealing the hierarchical organization of O_2_ diffusion pathways, this work introduces a generalizable, mutation-free approach for regulating gas access in metalloenzymes, offering conceptual and methodological guidance for the rational engineering of oxygen-tolerant biocatalysts.

## Conflict of Interest

The authors declare no conflict of interest.

## Data Availability Statement

All the data supporting these findings are available in the article and in the Supporting Information file.

## Author Contributions

X.S. conceived the study, performed all simulations and protein-design calculations, carried out the experiments, and wrote the original draft of the manuscript. WJ. L. co-conceived the study, executed some electrochmistry experiments, validated the resulting data, and critically reviewed the manuscript. WZ. L analyzed and visualized the experimental data and contributed to manuscript revision. H. L prepared samples and organized the raw data. Q.X. and LeYan.Z. participated in data analysis and manuscript revision. YL. F contributed in writing and editing. PY.J advised on protein-design strategies, optimized relevant protocols, and reviewed the manuscript. G.W. acquired funding, administered the project, supervised the research, and edited the manuscript. LiYun. Z. acquired funding, administered the project, supervised the entire study, and gave final approval of the manuscript.

All authors have read and agreed to the published version of the manuscript.

## Acknowledgments

This research was funded by the National Key R&D Program of China (grant numbers 2020YFA0907300), the National Natural Science Foundation of China (grant number 32170030).

## Entry for the Table of Contents

((Insert graphic for Table of Contents here (300 DPI resolution: up to 650×532 pixels for single-column format; up to 1358×295 pixels for double-column format). Please ensure your graphic is in **one** of the two following formats.))

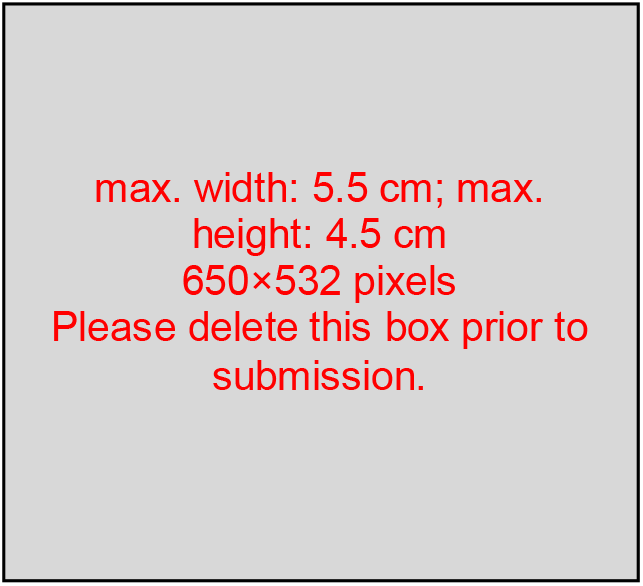

or

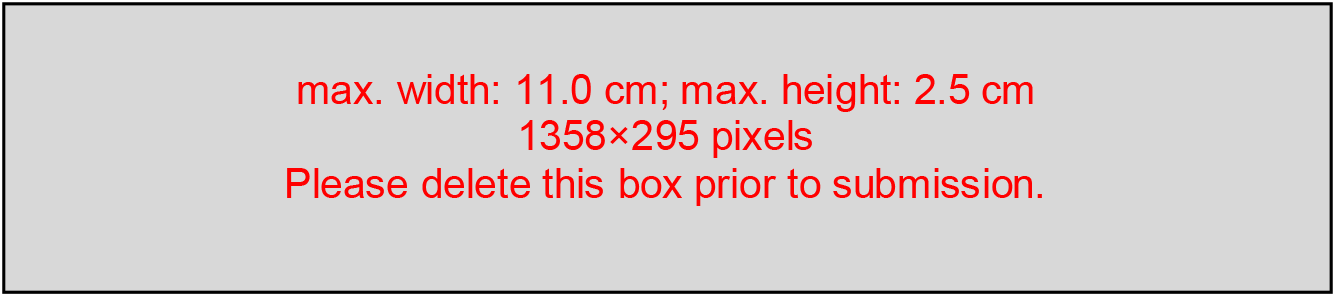

Insert text for Table of Contents here. ((Maximum 450 characters including spaces; the text should give readers a short preview of the main theme of the research and results, to attract their attention into reading the paper in full. Define acronyms, including those in the picture. The Table of Contents text should be different from the abstract.))

## Notes

### Competing Interest Statement

The authors have declared no competing interest.

## References

[1] Y. Shomura, M. Taketa, H. Nakashima, H. Tai, H. Nakagawa, Y. Ikeda, M. Ishii, Y. Igarashi, H. Nishihara, K. S. Yoon, S. Ogo, S. Hirota, Y. Higuchi, Science. 2017, 357, 928–932.

[2] B. L. Greene, C.-H. Wu, P. M. McTernan, M. W. W. Adams, R. B. Dyer, J. Am. Chem. Soc. 2015, 137, 4558–4566.

[3] M. W. Adams, L. E. Mortenson, J. S. Chen, Biochim. Biophys. Acta. 1980, 594, 105–176.

[4] F. Leroux, S. Dementin, B. Burlatt, L. Cournac, A. Volbeda, S. Champ, L. Martin, B. Guigliarelli, P. Bertrand, J. Fontecilla-Camps, M. Rousset, C. Leger, Proc. Natl. Acad. Sci. U. S. A. 2008, 105, 11188–11193.

[5] J. Kalms, A. Schmidt, S. Frielingsdorf, P. van der Linden, D. von Stetten, O. Lenz, P. Carpentier, P. Scheerer, Angew. Chem. Int. Ed. 2016, 55, 5586–5590.

[6] J. Kalms, A. Schmidt, S. Frielingsdorf, T. Utesch, G. Gotthard, D. von Stetten, P. van der Linden, A. Royant, M. A. Mroginski, P. Carpentier, O. Lenz, P. Scheerer, Proc. Natl. Acad. Sci. U. S. A. 2018, 115, E2229–E2237.

[7] Y. Montet, P. Amara, A. Volbeda, X. Vernede, E. C. Hatchikian, M. J. Field, M. Frey, J. C. Fontecilla-Camps, Nat. Struct. Biol. 1997, 4, 523–526.

[8] S. E. Beaton, R. M. Evans, A. J. Finney, C. M. Lamont, F. A. Armstrong, F. Sargent, S. B. Carr, Biochem. J. 2018, 475, 1353–1370.

[9] H. Ji, L. Wan, Y. Gao, P. Du, W. Li, H. Luo, J. Ning, Y. Zhao, H. Wang, L. Zhang, L. Zhang, J. Energy Chem. 2023, 85, 348–362.

[10] N. Plumere, O. Rudiger, A. A. Oughli, R. Williams, J. Vivekananthan, S. Poller, W. Schuhmann, W. Lubitz, Nat. Chem. 2014, 6, 822–827.

[11] A. A. Oughli, F. Conzuelo, M. Winkler, T. Happe, W. Lubitz, W. Schuhmann, O. Rüdiger, N. Plumeré, Angew. Chem. Int. Ed. 2015, 54, 12329–12333.

[12] J. Szczesny, J. A. Birrell, F. Conzuelo, W. Lubitz, A. Ruff, W. Schuhmann, Angew. Chem. Int. Ed. 2020, 59, 16506–16510.

[13] O. Ben-Zvi, I. Grinberg, A. A. Orr, D. Noy, P. Tamamis, I. Yacoby, L. Adler-Abramovich, ACS Nano. 2021, 15, 6530–6539.

[14] T. Li, Q. Jiang, J. Huang, C. M. Aitchison, F. Huang, M. Yang, G. F. Dykes, H.-L. He, Q. Wang, R. S. Sprick, A. I. Cooper, L.-N. Liu, Nat. Commun. 2020, 11, 5448.

[15] P. S. Huang, S. E. Boyken, D. Baker, Nature. 2016, 537, 320–327.

[16] J. Abramson, J. Adler, J. Dunger, R. Evans, T. Green, A. Pritzel, O. Ronneberger, L. Willmore, A. J. Ballard, J. Bambrick, S. W. Bodenstein, D. A. Evans, C. C. Hung, M. O’Neill, D. Reiman, K. Tunyasuvunakool, Z. Wu, A. Zemgulyte, E. Arvaniti, C. Beattie, O. Bertolli, A. Bridgland, A. Cherepanov, M. Congreve, A. I. Cowen-Rivers, A. Cowie, M. Figurnov, F. B. Fuchs, H. Gladman, R. Jain, Y. A. Khan, C. M. R. Low, K. Perlin, A. Potapenko, P. Savy, S. Singh, A. Stecula, A. Thillaisundaram, C. Tong, S. Yakneen, E. D. Zhong, M. Zielinski, A. Zidek, V. Bapst, P. Kohli, M. Jaderberg, D. Hassabis, J. M. Jumper, Nature. 2024, 630, 493–500.

[17] R. F. Alford, A. Leaver-Fay, J. R. Jeliazkov, M. J. O’Meara, F. P. DiMaio, H. Park, M. V. Shapovalov, P. D. Renfrew, V. K. Mulligan, K. Kappel, J. W. Labonte, M. S. Pacella, R. Bonneau, P. Bradley, R. L. Dunbrack, Jr., R. Das, D. Baker, B. Kuhlman, T. Kortemme, J. J. Gray, J. Chem. Theory. Comput. 2017, 13, 3031–3048.

[18] J. Dauparas, I. Anishchenko, N. Bennett, H. Bai, R. J. Ragotte, L. F. Milles, B. I. M. Wicky, A. Courbet, R. J. de Haas, N. Bethel, P. J. Y. Leung, T. F. Huddy, S. Pellock, D. Tischer, F. Chan, B. Koepnick, H. Nguyen, A. Kang, B. Sankaran, A. K. Bera, N. P. King, D. Baker, Science. 2022, 378, 49–56.

[19] J. L. Watson, D. Juergens, N. R. Bennett, B. L. Trippe, J. Yim, H. E. Eisenach, W. Ahern, A. J. Borst, R. J. Ragotte, L. F. Milles, B. I. M. Wicky, N. Hanikel, S. J. Pellock, A. Courbet, W. Sheffler, J. Wang, P. Venkatesh, I. Sappington, S. V. Torres, A. Lauko, V. De Bortoli, E. Mathieu, S. Ovchinnikov, R. Barzilay, T. S. Jaakkola, F. DiMaio, M. Baek, D. Baker, Nature. 2023, 620, 1089–1100.

[20] B. F. DL, A. P. Mikolajczyk, M. R. Carnes, W. Sharp, E. Revellame, R. Hernandez, W. E. Holmes, M. E. Zappi, Biotechniques. 2024, 76, 14–26.

[21] K. H. Sumida, R. Nunez-Franco, I. Kalvet, S. J. Pellock, B. I. M. Wicky, L. F. Milles, J. Dauparas, J. Wang, Y. Kipnis, N. Jameson, A. Kang, J. De La Cruz, B. Sankaran, A. K. Bera, G. Jimenez-Oses, D. Baker, J. Am. Chem. Soc. 2024, 146, 2054–2061.

[22] N. R. Bennett, B. Coventry, I. Goreshnik, B. Huang, A. Allen, D. Vafeados, Y. P. Peng, J. Dauparas, M. Baek, L. Stewart, F. DiMaio, S. De Munck, S. N. Savvides, D. Baker, Nat. Commun. 2023, 14, 2625.

[23] N. R. Bennett, J. L. Watson, R. J. Ragotte, A. J. Borst, D. L. See, C. Weidle, R. Biswas, Y. Yu, E. L. Shrock, R. Ault, P. J. Y. Leung, B. Huang, I. Goreshnik, J. Tam, K. D. Carr, B. Singer, C. Criswell, B. I. M. Wicky, D. Vafeados, M. G. Sanchez, H. M. Kim, S. V. Torres, S. Chan, S. M. Sun, T. Spear, Y. Sun, K. O’Reilly, J. M. Maris, N. G. Sgourakis, R. A. Melnyk, C. C. Liu, D. Baker, BioRxiv. 2025, 2024.2003.2014.585103.

[24] E. Chovancova, A. Pavelka, P. Benes, O. Strnad, J. Brezovsky, B. Kozlikova, A. Gora, V. Sustr, M. Klvana, P. Medek, L. Biedermannova, J. Sochor, J. Damborsky, PLoS Comput. Biol. 2012, 8, e1002708.

[25] D. Sehnal, R. Svobodova Varekova, K. Berka, L. Pravda, V. Navratilova, P. Banas, C. M. Ionescu, M. Otyepka, J. Koca, J. Cheminform. 2013, 5, 39.

[26] V. H. Teixeira, A. M. Baptista, C. M. Soares, Biophys. J. 2006, 91, 2035–2045.

[27] S. Zacarias, A. Temporao, P. Carpentier, P. van der Linden, I. A. C. Pereira, P. M. Matias, J. Biol. Inorg. Chem. 2020, 25, 863–874.

[28] J. Jumper, R. Evans, A. Pritzel, T. Green, M. Figurnov, O. Ronneberger, K. Tunyasuvunakool, R. Bates, A. Zidek, A. Potapenko, A. Bridgland, C. Meyer, S. A. A. Kohl, A. J. Ballard, A. Cowie, B. Romera-Paredes, S. Nikolov, R. Jain, J. Adler, T. Back, S. Petersen, D. Reiman, E. Clancy, M. Zielinski, M. Steinegger, M. Pacholska, T. Berghammer, S. Bodenstein, D. Silver, O. Vinyals, A. W. Senior, K. Kavukcuoglu, P. Kohli, D. Hassabis, Nature. 2021, 596, 583–589.

[29] C. Han, Q. Liu, X. Luo, J. Zhao, Z. Zhang, J. He, F. Ge, W. Ding, Z. Luo, C. Jia, L. Zhang, Biosens. Bioelectron. 2024, 116313.

[30] L. Zhang, X. Fang, X. Liu, H. Ou, H. Zhang, J. Wang, Q. Li, H. Cheng, W. Zhang, Z. Luo, Chem. Commun. 2020, 56, 10235–10238.

[31] F. A. Armstrong, B. Cheng, R. A. Herold, C. F. Megarity, B. Siritanaratkul, Chem. Rev. 2023, 123, 5421–5458.

[32] F. A. Armstrong, R. M. Evans, S. V. Hexter, B. J. Murphy, M. M. Roessler, P. Wulff, Acc. Chem. Res. 2016, 49, 884–892.

[33] J. Kalms, A. Schmidt, S. Frielingsdorf, T. Utesch, G. Gotthard, D. von Stetten, P. van der Linden, A. Royant, M. A. Mroginski, P. Carpentier, O. Lenz, P. Scheerer, Proc. Natl. Acad. Sci. U. S. A. 2018, 115, E2229–E2237.

[34] M. J. Lukey, A. Parkin, M. M. Roessler, B. J. Murphy, J. Harmer, T. Palmer, F. Sargent, F. A. Armstrong, J. Biol. Chem. 2010, 285, 3928–3938.

[35] R. M. Evans, S. E. Beaton, P. R. Macia, Y. Pang, K. L. Wong, L. Kertess, W. K. Myers, R. Bjornsson, P. A. Ash, K. A. Vincent, S. B. Carr, F. A. Armstrong, Chem. Sci. 2023, 14, 8531–8551.

[36] M. del Barrio, C. Guendon, A. Kpebe, C. Baffert, V. Fourmond, M. Brugna, C. Leger, ACS Catal. 2019, 9, 4084–4088.

[37] T. Buhrke, O. Lenz, N. Krauss, B. Friedrich, The Journal of biological chemistry. 2005, 280, 23791–23796.

[38] E. Garcin, X. Vernede, E. C. Hatchikian, A. Volbeda, M. Frey, J. C. Fontecilla-Camps, Structure. 1999, 7, 557–566.

[39] A. Parkin, G. Goldet, C. Cavazza, J. C. Fontecilla-Camps, F. A. Armstrong, J. Am. Chem. Soc. 2008, 130, 13410–13416.

[40] M. E. Pandelia, V. Fourmond, P. Tron-Infossi, E. Lojou, P. Bertrand, C. Léger, M. T. Giudici-Orticoni, W. Lubitz, J. Am. Chem. Soc. 2010, 132, 6991–7004.

